# *N*-Glycopedia: Libraries for Native *N*-glycan Structural Analysis

**DOI:** 10.1101/2025.06.09.658590

**Authors:** Christopher Ashwood, Richard D Cummings

## Abstract

Studies on glycans of glycoproteins are hampered by the lack of standards that reflect the wide diversity in structure typically observed. To this end we have exploited a large library of *N*-glycan standards comprised of a unique collection of 226 *N*-glycans including oligomannose, hybrid, and complex-type. We generated a method employing porous graphitized carbon (PGC) and liquid chromatography mass spectrometry (PGC-LC-MS), which can provide a high degree of resolution of underivatized *N*-glycan structures. Chromatogram libraries arising from these studies include retention time data, diagnostic fragments, and validated structural assignments, providing a robust platform for both targeted and discovery-based glycomics. We refer to this as an *N*-glycopedia, the first type of resource in which researchers can compare this collective data to *N*-glycans under study and overcome the limitations of only having compositional data and predicted structures. The technology is easily expandable to include additional *N*-glycans as new standards become available.

## Introduction

Protein glycosylation represents one of the most complex and crucial post-translational modifications in biology, playing pivotal roles in cellular recognition, protein folding, and immune responses^1^. The diversity of glycan structures arises from interactions between species-specific enzymatic machinery, cellular compartmentalization, and protein-specific factors^2–4^. This complexity presents a significant analytical challenge, particularly in distinguishing structural isomers that share identical masses but exhibit distinct biological functions^5,6^.

Traditional approaches to glycan analysis have relied heavily on chemical derivatization, specifically reduction, permethylation, and reducing-end tags, to enhance detection sensitivity and structural characterization^7^. However, these modifications can introduce analytical artifacts and increase sample preparation complexity^8,9^. Recent advances in analytical technologies, particularly in liquid chromatography coupled to mass spectrometry (LC-MS), have opened new possibilities for analyzing native glycans. Porous graphitized carbon (PGC) liquid chromatography has emerged as a powerful tool for separating glycan isomers, while mass spectrometry provides detailed structural information through diagnostic fragmentation patterns^10,11^.

Triple quadrupole mass spectrometers have become the gold standard for clinical diagnostics and targeted quantification due to their exceptional sensitivity, specificity, and robust performance^12^. Selected reaction monitoring (SRM) mode, where specific precursor-to-product ion transitions are monitored, enables precise quantification even in complex biological matrices due to the specificity enabled by precursor ion mass, product ion mass, collision energy (CE), and retention time (RT) variables. This targeted approach has revolutionized clinical assays, from small molecule drug monitoring to protein biomarker detection, offering unparalleled reproducibility and analytical precision^13^. However, in glycomics, the application of triple quadrupole MS has been limited by the requirement for pre-defined fragmentation patterns and purified standards for method development and validation through quantifier-qualifier ion ratio assessment. Flowers *et. al.^14^* have an established targeted *O*-glycan SRM method offering substantial insight in the clinical osteoarthritis area, but an equivalent *N*-glycan assay does not exist despite its potential for clinical impact.

Here we present *N*-glycopedia, a comprehensive analytical framework for native *N*-glycan structure analysis built upon the systematic characterization of 226 purified glycan standards using triple quadrupole MS. We developed an optimized PGC-LC-MS method capable of resolving unmodified *N*-glycans with unprecedented separation of structural isomers. The resulting chromatogram libraries incorporate RT data, diagnostic fragment ions, and validated structural assignments, providing a robust platform for both targeted and discovery-based glycomics.

## Results

### A facile, targeted N-glycan assay is enabled by non-reduced glycan analysis

Purified, unmodified *N*-glycan standards from commercial sources were initially profiled with SUGA, a compositional analysis method^15^, to assess non-isomeric purity and define the product ion spectra generated by each standard (**Figure 1A**). With a total of 226 *N*-glycan standards that met purity requirements (>95% pure), this initial screen was rapidly completed in 12 hours. To enable RT normalisation across each standard and exploit the RT dimension for isomer discrimination, each standard plate was dried and standards were resuspended in an aqueous solution containing partially hydrolysed dextran.

**Figure 1.**
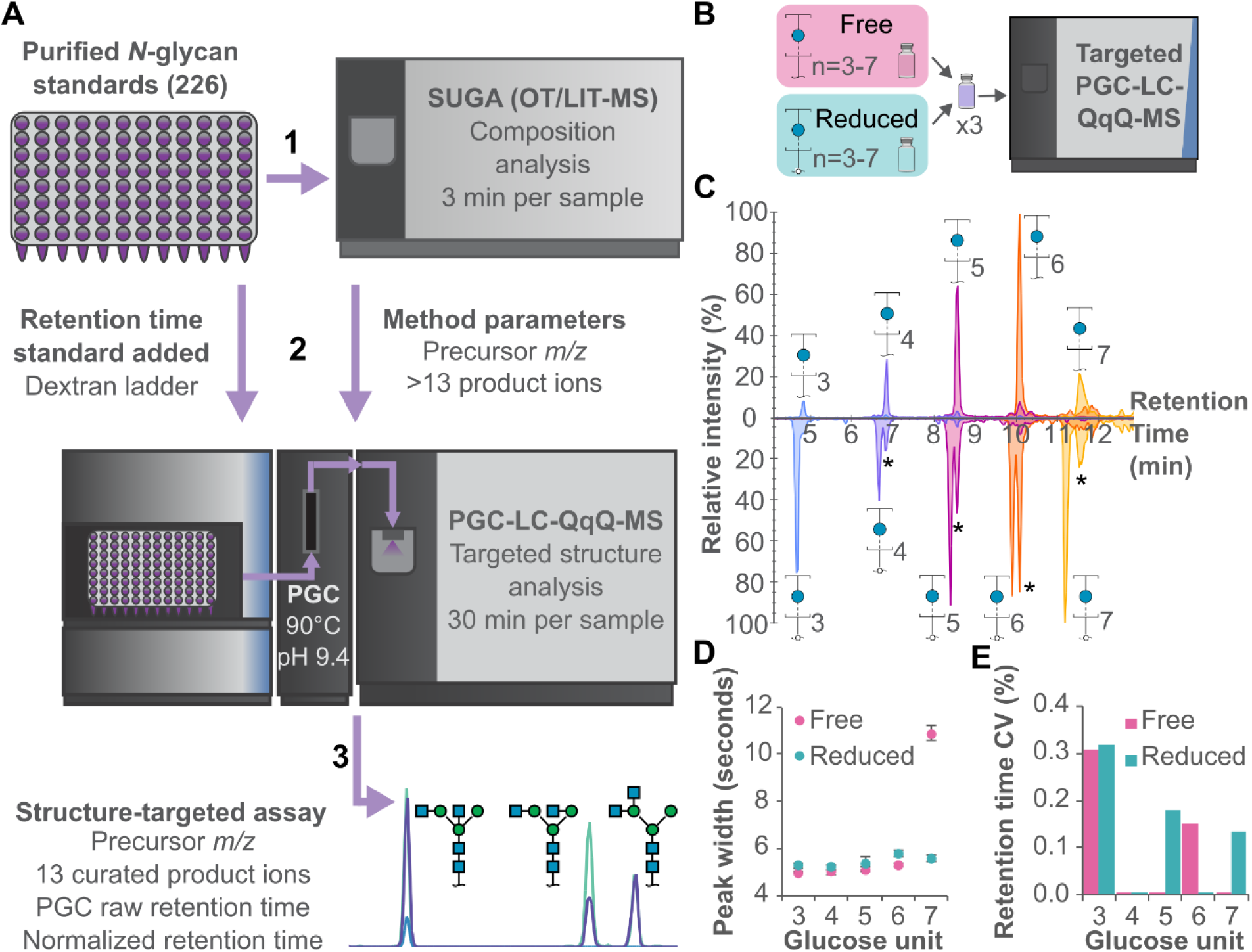
Advancements in non-reduced glycan analysis enable the development of a targeted *N*-glycan assay. **(A)** Data acquisition workflow for constructing an *N*-glycopedia of pure *N*-glycan structures. **(B)** Comparison of reduced and non-reduced glycans demonstrating equivalent performance by native PGC-LC-MS. **(C)** Reduced and non-reduced dextran ladder subunits produce equivalent chromatograms (* indicates isotopic interference from non-reduced dextran ladder). **(D)** Peak widths are consistent across dextran ladder formats, with the exception of GU7. **(E)** RTs are consistent and equivalent between dextran ladder formats.

Subsequently, the standards were then analysed by a targeted, standard-specific PGC-LC-QqQ-MS SRM assay, based on the observed product ions in the SUGA acquisition. The incorporation of PGC-LC enables isomer-specific separation, assessing structural purity which is not determined by SUGA. Additionally, the SRM assay measures each product ion at nine different collision energies (CEs), achieving structure-specific optimisation of each product ion. As a result of the overall workflow, we developed a structure-targeted assay composed of a precursor *m/z*, 13 product ions with optimal CEs, raw PGC RT, and a normalised PGC RT for 183 unique structures corresponding to 91 glycan compositions (**Figures S1 and S2**).

Typically, glycan reduction is an essential sample preparation step prior to PGC separation of glycans due to its high resolving power separating alpha and beta anomers which are artifactual and irrelevant for glycan analysis^16^. Using our unique combination of PGC column temperature, pH, and mobile phases, we eliminate the anomeric effect on PGC. A mixture of reduced and non-reduced dextran ladder run with a targeted method demonstrates similar peak widths and RTs regardless of the reduction state (**Figure 1B and C**). We observed a doubling of peak width at 7 GU however all other parameters, including RT variation were similar or identical (**Figure 1D and E**). With column temperature optimisation, larger non-reduced dextran ladder subunits were observed to be more sensitive to peak broadening, and temperatures of 70°C and 90°C observed as suitable for generating tight peaks (**Figure S3**).

### Benchmarking native PGC-LC by diverse standards verifies structural separation

The vendor diversity of glycan standards used to construct *N*-glycopedia enable a larger, composite library with few redundancies (**Figure 2A**). One such redundancy, G36191CD, a bi-antennary galactose-capped *N*-glycan, was produced by all five vendors and serves as a way to assess inter-vendor agreement despite variations in synthesis route. A mixture of each vendor’s glycan standard was made and analysed by our targeted method, yielding a single peak with no isomeric impurities observed (**Figure 2B**). Additional redundant structures were analysed by this same approach, with all seven cross-validations generating a single, dominant peak (**Figure S4**). This demonstrates agreement and consistency across glycoscience vendors, especially in the case of stable isotope labelled glycan co-elution as generated by Vendor C.

**Figure 2.**
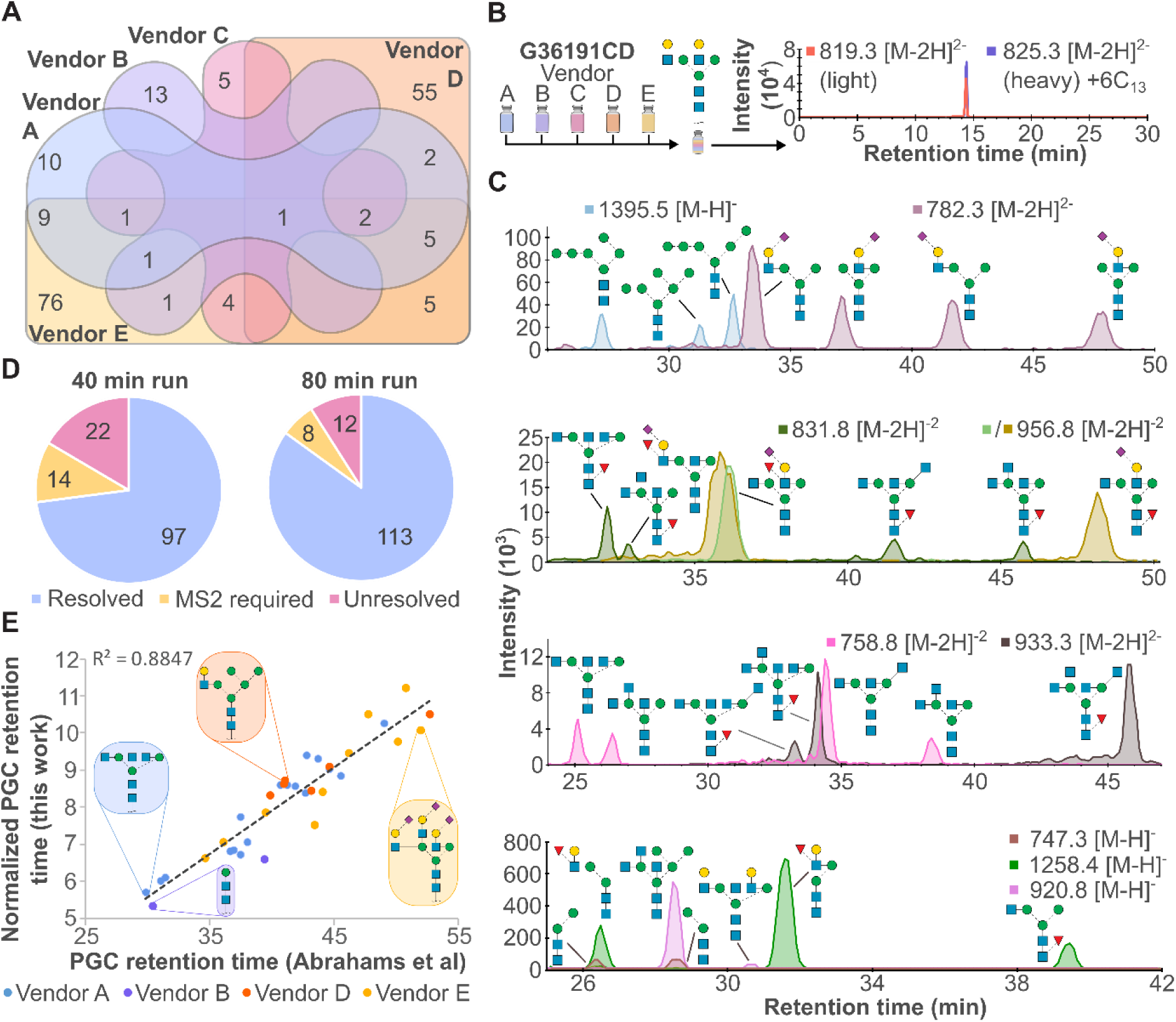
Benchmarking of glycan standards by native PGC-LC reveals consistent performance across a broad structural repertoire. **(A)** Venn diagram showing overlap in structural identity across different vendors. **(B)** All vendors produce the same pure structure, regardless of synthetic route. **(C)** Native PGC-LC-MS resolves challenging isomeric mixtures, including mannosylation, sialylation, fucosylation, galactosylation, and GlcNAcylation. **(D)** PGC-LC gradient optimization minimizes reliance on MS2 for structural identification. **(E)** Consistency of native PGC-LC values for *N*-glycan standards compared to published data from Abrahams *et. al.^10^*

Following analysis of each individual standard, our separation approaches were benchmarked using isomeric mixtures including mannose, sialic acid, fucose, galactose, and *N*-acetylglucosamine (GlcNAc) linkage and topology isomers. As shown across **Figure 2C**, these isomers were all baseline resolved by LC except for two structures with different arm positions but the same sialic acid and fucose linkages (956.8 *m/z*). This showcases the high resolving power of PGC and the ability to resolve non-reduced glycan isomers when anomers are collapsed into a single peak. Gradient length was assessed and a doubling of acquisition time resulted in an 18% improvement in isomers being resolved (**Figure 2D**, 72% vs 85% resolved).

Finally, as PGC RT is useful dimension for structural elucidation and verification, glycan RT libraries have been constructed such as GlycoStore^17^. Using glycan structures shared between our library and their assessed structures, we observe a strong positive relationship between our normalised PGC RTs and the raw PGC RTs provided by Abrahams *et. al.^10^* (**Figure 2E**). The largest disagreements occurred for paucimannose and highly sialylated *N*-glycans. This strong correlation is beneficial in two ways: firstly, it demonstrates non-reduced glycans have near identical RT patterns as reduced glycans, and secondly, our glycan library can expand upon this existing resource.

### Structure-specific fragmentation optimisation enables isomer discrimination

The structural specificity of product ions serves as the primary method to specifically and sensitively measure each glycan however they require experimental identification and optimisation. After product ion optimisation, we found that total intensity for each standard’s set of product ions could be predicted with reasonable accuracy based on the positive relationship between optimal CE and precursor *m/z* (**Figure 3A**). At the individual product ion level, however, there was a poor correlation between product m/z and optimal CE, demonstrating that each product ion should be experimentally optimised (**Figure 3B**). This observation contrasts SRM-based proteomics, where optimal CE could be confidently estimated based on product ion *m/z^18^*, likely owing to the linear nature of peptides compared to non-linear glycans. This unpredictability could also reflect competing fragmentation mechanisms within a glycan, which could be exploited to discriminate structural isomers.

**Figure 3.**
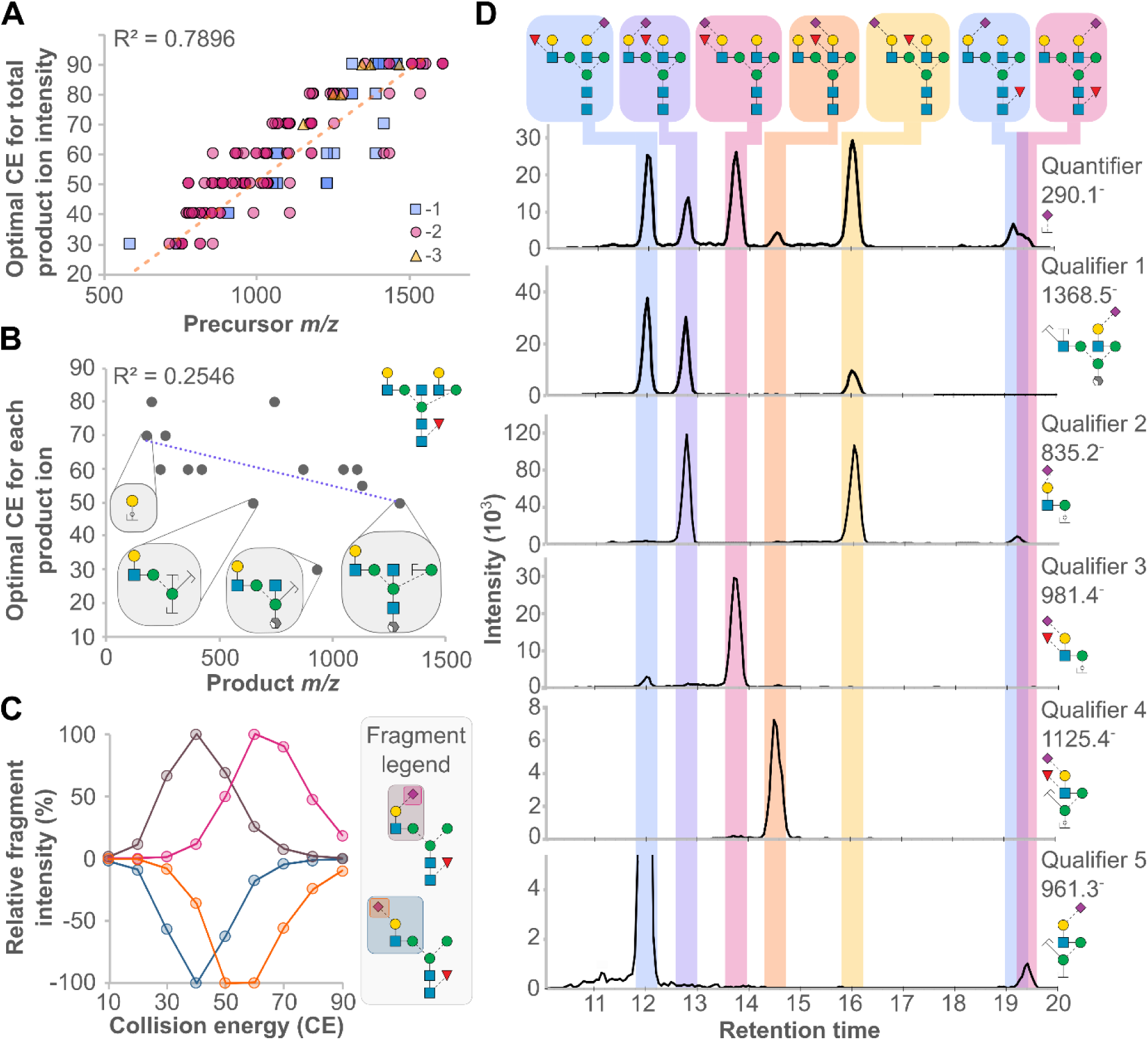
Collision energy (CE) optimization reveals structural dependencies in fragment ion intensities for isomer discrimination. **(A)** The optimal CE for achieving maximum total product ion intensity can be predicted based on precursor *m/z*. **(B)** The optimal CE for individual product ions shows a weak correlation with product *m/z*. **(C)** Distinct CE profiles are observed for different monosaccharide linkages. **(D)** Glycan isomer discrimination is facilitated by glycotope-specific product ion combinations

Glycan motif-specific fragmentation patterns were evaluated and linkage-dependent CE differences were observed and exploited, enabling discrimination based on precursor-product ion pairs alone. Two structures, differing only by their sialic acid linkage, had identical CE plots for a product ion representing the cleavage of the same glycosidic bond (**Figure 3C**, α1,3 mannose). This is contrasted by their sialic acid fragments featuring different CE plots, with α2,3 linked sialic acid requiring 10 CE units more to fragment abundantly compared to its α2,6 linked counterpart. Although equivalently intense at 60 CE, specificity could be enhanced by using 50 or 70 CE for α2,3 and α2,6 sialic acid linkage, respectively. Isomer discrimination for all structures in the glycan library was established based on these principles as well as motif-specific fragments.

Seven structural isomers, varying in fucose and sialic acid position and linkage, were characterised by their unique product ion profiles as a proof of concept for the overall assay (**Figure 3D**). The quantifier for each of the structures was a shared, abundant fragment, 290.1 *m/z*, which inherently was not used as a structure-specific ion. Qualifier 1 is a product ion specific for 3/7 of the structures featuring sialic acid on one arm, and fucose attached to the other. Qualifier 2 is specific for structures with sialylated, non-fucosylated α1,3 arms.

Structures with antennae decorated with both fucose and sialic acid do not produce these products ions, requiring qualifiers 3 and 4 to differentiate α1,3-arm modification and α1,6-arm modification, respectively. Finally, qualifier 5 is used to specifically measure non-fucosylated 6-arm modification with α2,6 sialic acid. In this example, this series of qualifier ions enables each motif in each glycan structure to be simultaneously measured, and finally combined as an ultimate representation of a given structure. This approach contrasts the diagnostic ions previously defined and used frequently in untargeted mass spectrometry experiments^11,19^, as qualifier ions for an SRM assay are defined before acquisition.

### N-glycopedia enables reproducible targeted glycan structure assays

Using the principles described through this work (standard characterization, native isomer LC separation, and structure-specific product ions with defined collision energy), a single PGC-LC-QqQ-MS *N*-glycan assay was defined to specifically measure 183 unique, mammalian glycan structures with 431 unscheduled transitions. Unlike typical SRM assays, glycan transitions were unscheduled to enable the method to be used across a wide range of chromatographic setups, and flexibility to quantify glycan structures not originally used to build the assay (**Figure S5**) across 93 glycan compositions (**Figure S2**). Data analysis is expedited using a spectral library comprised of a GlyTouCan accession representing the structure measured, the expected relative intensities for each structural qualifier and their normalised RT values (**Figure 4A**). These analyses were performed using Skyline, a transparent, vendor-neutral software platform. This final assay could then be applied to any *N*-glycosylated sample type.

**Figure 4.**
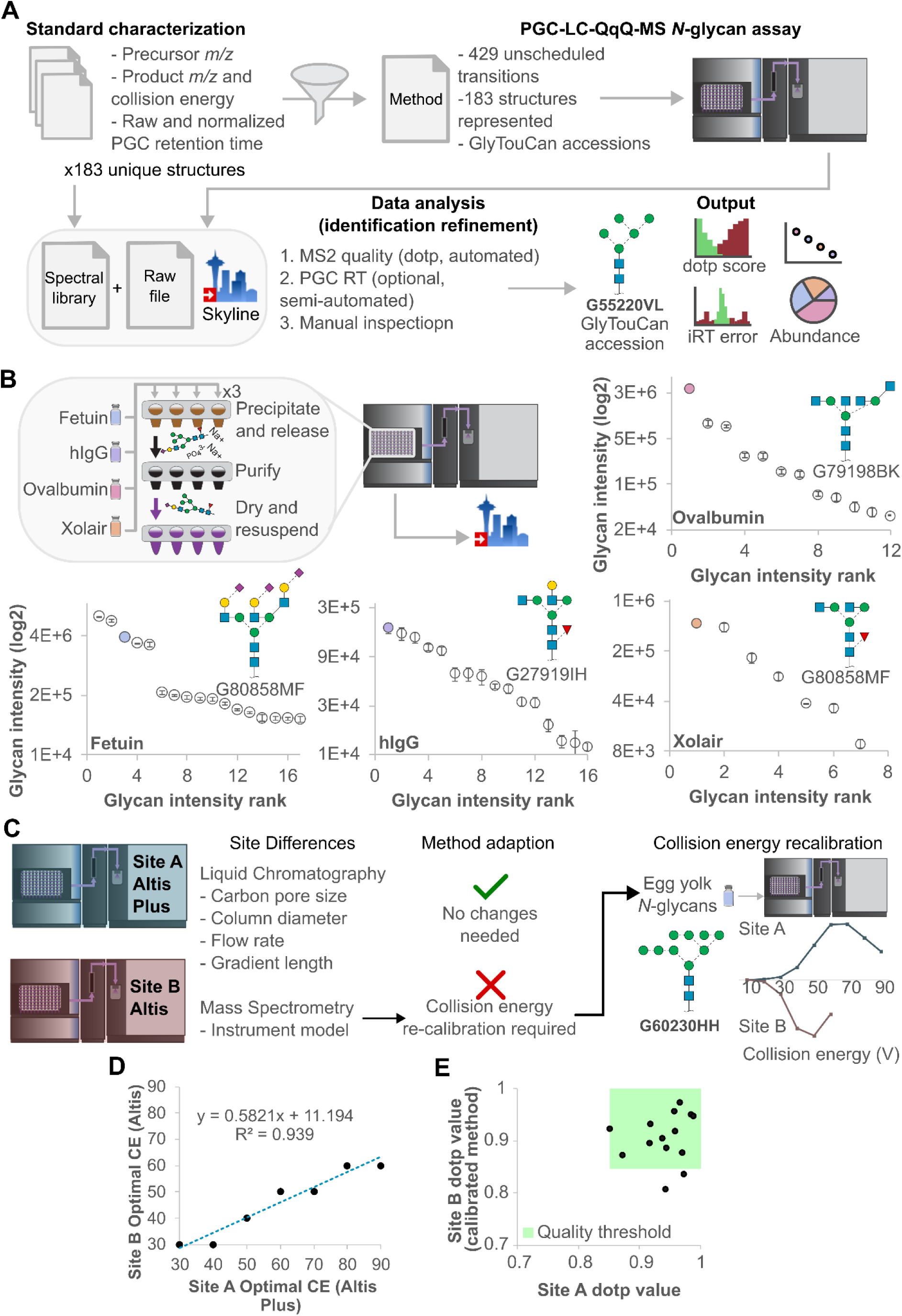
*N*-glycopedia enables reproducible targeted glycan structure analysis across glycoproteins and laboratories. **(A)** Chromotogram libraries are filtered to define an *N*-glycan structure assay with rapid data analysis and result contextualisation. **(B)** Structural quantitation of *N*-glycans released from purified glycoproteins demonstrates method repeatability. **(C)** Collision energy re-calibration is the only method adaption step required for method translation across laboratories. **(D)** Optimal collision energy plot for method adaption. **(E)** Quality *N*-glycan identifications for bovine fetuin are consistent across sites.

The sample preparation from our SUGA workflow, which rapidly releases and purifies non-reduced glycans^15^, was applied to a set of glycoproteins in triplicate to demonstrate our SRM assay generates repeatable, structure-specific quantitation with anomers collapsed into a single peak (**Figure 4B**). *N*-glycan profiles observed were consistent with previously published data, covered all expected structures of these model glycoproteins, and total variation for all structures was <20% co-efficient of variation (CV) over 2 orders of magnitude dynamic range for all but one structure, consistent with a clinical assay requirement for MS assays^20^.

The usability and translatability of this assay to other labs is key and steps required to translate our assay to a different LC-MS system. Due to the unscheduled nature of our transitions, site differences in liquid chromatographic setup did not require method adaption, however CE recalibration was required to account for a different mass spectrometer model (**Figure 4C**). In this example, we profiled *N*-glycans from a cheap, commercially available source, egg yolk, specifically chosen due to its lack of isomers, and acquired a CE profile on the Site B mass spectrometer. Comparing the CE plot at Site B to the purified *N*-glycan standard CE plot at site A, we observed a significant shift in the apex CE value and hypothesised this could be used to recalibrate the SRM assay CE values from Site A.

Optimal CE for 20 product ions corresponding to seven observed *N*-glycans in egg yolk was compared between sites. We observed a strong linear correlation between Site A and Site B optimal CE values, and used the resulting equation to harmonize the SRM assay CE values (**Figure 4D**). After CE recalibration, the egg yolk N-glycan sample data was re-acquired and an approximately 5% improvement in product ion dotp score was observed (**Figure S6**). The effectiveness of harmonization between sites was evaluated by bovine fetuin *N*-glycans released and analysed at both sites. Using the *N*-glycopedia spectral library in Skyline, we observed all but two *N*-glycans reaching sufficient quality thresholds for classification as confident identifications (**Figure 4E**).

*N*-glycopedia was then applied to study samples with expected differences in host glycosylation, glycoprotein site-specific accessibility, and defects in glycosylation pathways through SARS-CoV2 spike glycoproteins, human immunoglobulins, and mutant CHO cell line with defective *N*-glycosylation. SARS-CoV2 spike proteins expressed by Human (HEK293), Hamster (CHO), and Insect (CF9) cell lines, and often used as models of in-vivo spike glycoproteins, yielded distinct *N*-glycan profiles caused by differences in their host machinery (**Figure 5A**). HEK293 spike glycoprotein featured the most diverse glycan profile with predominantly complex bi-antennary glycans but also featuring a significant proportion of oligomannose glycans, corresponding well with previously published glycoproteomic studies. CHO spike glycoprotein was more homogenous, with a single α2,3 bi-antennary *N*-glycan making up more than 30% of the observed glycan peak area, and other glycan classes represented 26% of observed glycans. Lastly, the SF9-expressed spike glycoprotein was ∼70% of a core-fucosylated paucimannose structure with minor oligomannose and complex mono-antennary classes, suitable for our assay despite expression in an insect host. The versatility of *N*-glycopedia for characterizing diverse glycoproteins was further demonstrated by its application to human immunoglobulin glycoproteins, revealing critical insights into their structure-function relationships.

**Figure 5.**
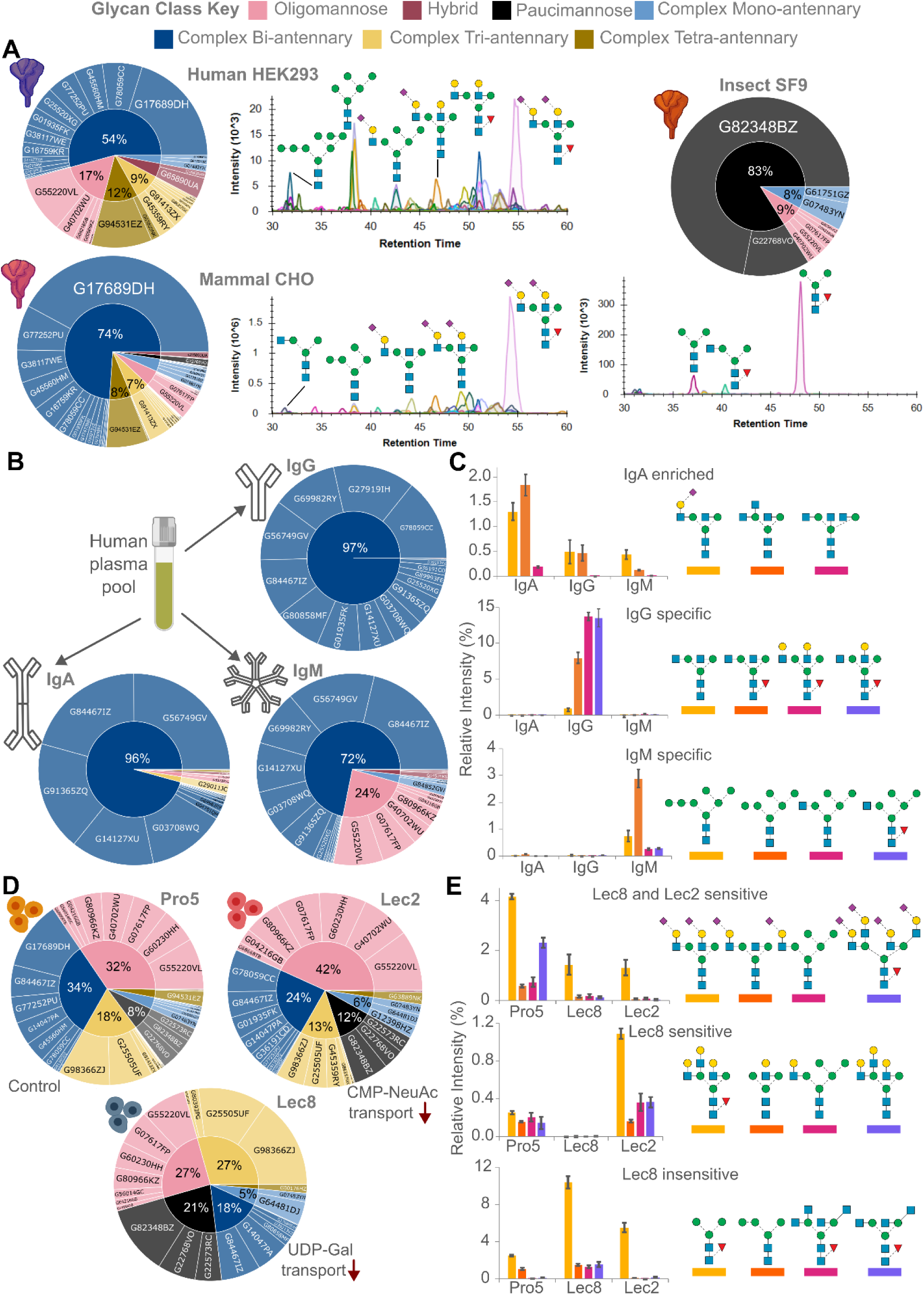
Targeted quantitative data across recombinant protein expression systems, purified human immunoglobulins, and perturbated cell lines reveals structure-specific expression. **(A)** Technical triplicate N-glycan profiles from SARS-CoV-2 spike glycoprotein expressed in SF9 insect cells, CHO mammalian cells, and HEK293 cells. Discrimination between IgA, IgG, and IgM from a human plasma pool using *N*-glycan **(B)** class profiles and **(C)** structure abundances in technical triplicate. Restriction of nucleotide sugar transport into the Golgi results in profound changes in N-glycan **(D)** class profiles and **(E)** structure abundances in technical triplicate.

Quantitative comparisons performed on the *N*-glycans released in triplicate and measured by SRM for IgA, IgG, and IgM demonstrate the effectiveness of our method for sensitive glycan detection. In addition to class differences between those immunoglobulins, significant structure-specific differences were observed that enabled differentiation between the three glycoproteins (**Figure 5B**). IgA lacked specific *N*-glycans, rather three *N*-glycans were at least 2-fold more abundant in IgA, with the last, a bisected bi-antennary *N*-glycan was 10-fold more intense (**Figure 5C**). Four *N*-glycans were at least 10-fold more abundant in IgG: bi-antennary terminated with galactose or GlcNAc. Conversely, oligomannose and hybrid *N*-glycans were significantly >10-fold more abundant in IgM. Our *N*-glycopedia methodology thus successfully detected structural variations across all immunoglobulin classes, while its quantitative precision revealed significant differences in glycan distribution that align with the distinct functional roles of these critical immune proteins^21^.

Contrasting the purified glycoproteins characterised, we then characterised Pro5, Lec2, and Lec8 cell lines which are the control, sialic acid deficient (via low CMP-NeuAc in the Golgi), and galactose deficient (via low UDP-Gal in the Golgi), respectively^22^ (**Figure 5D**). Sialylation was significantly more abundant in the Pro5 control cell line compared to the Lec8 and Lec2 cell lines due to either decreased Golgi CMP-Sia, or decreased *N*-glycan galactosylation, a requirement for subsequent sialylation (**Figure 5E**). Galactosylated tri-antennary, mono-antennary, and hybrid *N*-glycans were significantly increased in Pro5 and Lec2 cell lines, with more in the Lec2 cell line due to the galactosylated substrate being unable to be sialylated. Finally, paucimannosidic, hybrid, and GlcNAc-capped *N*-glycans were significantly more abundant in the Lec8 cell line.

## Discussion

The development of *N*-glycopedia represents a significant advancement in glycan analysis, offering unprecedented structural specificity without the need for glycan reduction. By combining native PGC-LC separation with optimized collision energy profiles for structure-specific product ions, this methodology enables reliable identification and quantification of 183 unique *N*-glycan structures. The strong correlation between our normalized RTs and previously published data validates our approach while demonstrating that non-reduced glycans exhibit retention patterns nearly identical to their reduced counterparts. This finding not only simplifies sample preparation but also allows integration with existing glycan libraries, effectively building upon rather than replacing previous glycoscience knowledge.

Our application of *N*-glycopedia across diverse biological systems demonstrates its versatility and biological relevance. The distinct *N*-glycan profiles observed in SARS-CoV-2 spike glycoproteins expressed in different cell lines highlight the significant impact of host glycosylation machinery on recombinant protein production, with HEK293-expressed spike showing the most diverse profile compared to CHO and SF9. Similarly, the quantitative differences detected between immunoglobulin classes (IgA, IgG, and IgM) align with their distinct functional roles in the immune system. Perhaps most compelling is the methodology’s ability to precisely characterize glycosylation defects in Lec2 and Lec8 cell lines, providing direct evidence of pathway-specific alterations in glycan processing that correspond with known deficiencies in nucleotide sugar availability.

The robust inter-laboratory transferability of *N*-glycopedia addresses a critical need in glycan analysis for standardized, reproducible methods. There have been prior studies aimed at generating libraries of N-glycans^23–25^), but several methods rely on MALDI-TOF and use virtual libraries for compositional analyses and computational predictions, or have limited number of synthetic N-glycan standards, rather than large libraries of *N*-glycans and a novel technology to separate isomers.

In the methods described here, with only collision energy recalibration required between different MS platforms, this targeted approach offers a practical solution for consistent glycan structure identification across research sites. The comprehensive spectral library, combined with transparent data analysis through vendor-neutral software, facilitates broad adoption of this methodology in both research and clinical settings. As glycan structure increasingly becomes recognized as a critical determinant of biological function, *N*-glycopedia provides a powerful tool for precise glycan characterization that should accelerate discoveries across glycobiology, immunology, and biotherapeutic development.

Our approach addresses several key challenges in the field. First, it detects phylogenetic diversity of glycan structures present across protein expression hosts, encompassing variations observed across different phyla and orders, from insects to primates. Second, it captures the complexity introduced by cell-specific enzymatic machinery, including nucleotide sugar transporters, and glycosyltransferases. Third, it reflects the influence of site-specific accessibility on glycan processing, providing insights into protein-specific glycosylation patterns.

The *N*-glycopedia framework represents a significant advance over existing methods by eliminating the need for glycan reduction while maintaining high analytical performance. This approach not only simplifies sample preparation but also preserves valuable structural information that might be lost during derivatization. The comprehensive nature of our chromatogram libraries, combined with detailed fragmentation patterns, enables reliable identification of glycan structures without requiring individual reference standards for each analysis. By leveraging the sensitivity and specificity of triple quadrupole MS, our method achieves quantitative precision comparable to clinical assays, making it suitable for routine glycan monitoring in both research and clinical settings.

Furthermore, we demonstrate the utility of *N*-glycopedia through extensive benchmarking against existing methods and its application to diverse biological samples. Our framework enables both targeted quantification and untargeted discovery of *N*-glycans, providing researchers with a versatile tool for glycomics research. The standardized protocols and detailed documentation ensure reproducibility across different laboratories and instrumental platforms, addressing a critical need in the glycomics community for robust, validated analytical methods.

This work establishes a foundation for systematic native *N*-glycan analysis, offering the glycobiology community a powerful resource for advancing our understanding of protein glycosylation in health and disease. The principles and methodologies presented here can be extended to other classes of glycans, potentially transforming our ability to characterize the complex glycomes of biological systems while maintaining the analytical rigor required for clinical applications.

## Online methods

### Cell Culture and Lysis

Cell lines (Pro5, Lec2, and Lec8^22^) were grown according to the supplier’s instructions (American Type Culture Collection – ATCC in Manassas, VA), in media supplemented with 10% fetal bovine serum. Six cell culture replicates were grown to approximately a 50% density in 6-well plate and allowed to grow for 24–48 h at 37°C until confluent (∼10^5^ cells per well). The medium was aspirated and each well was washed four to five times with PBS. The cell pellets were collected and frozen for downstream preparation. Frozen cells were lysed in 10% SDS via probe sonication (3x 10 seconds on, 10 seconds off).

### Glycoprotein Reduction, Alkylation, and Glycan Release

Human immunoglobulins (IgG, IgM, IgA) were purchased from Athens Research & Technology (Athens, GA). SARS-CoV2 spike proteins were produced and sourced from BEI Resources (Biodefense and Emerging Infections Research Resources Repository) - see https://www.beiresources.org/Home.aspx. Purified glycoproteins for initial profiling were purchased from Sigma-Aldrich (St. Louis, MO). Xolair glycoprotein, which is a recombinant IgG, was a kind gift from Dr. Robert Anthony, Harvard Medical School. The purified and cell lysate proteins were reduced with 5 mM dithiothreitol for 30 min at 55 °C, alkylated with 10 mM iodoacetamide in the dark for 30 min at room temperature, and quenched with an additional 5 mM dithiothreitol for 15 min at room temperature. Glycans were released from glycoproteins and purified for LC-MS analysis as described in Ashwood *et al^15^*. In brief, proteins were precipitated with methanol and phosphoric acid, and purified via silica column. Glycans were released with PNGase-F (New England Biolabs, Ipswich, MA), purified by adsorption to PGC solid phase extraction, and then dried prior to LC-MS analysis.

### Purified Glycan Standard Preparation for LC and MS Benchmarking/Calibration

Internal standards for RT alignment and external glycan standards for MS CE calibration were performed using starting material of dextran, and dried egg yolk protein, respectively. Dextran ladder was prepared as described in Ashwood *et al^26^*. Egg yolk glycans were released as described above for purified glycoprotein release and purification.

### Purified N-glycan Standards

*N*-glycan standards were sourced from five vendors (Dextra Laboratories Ltd (Reading, UK), Omicron Biochemicals Inc. (South Bend, IN), Glycobia Inc.^6^ (Ithaca, NY), Chemily LLC^27^ (Peachtree Corners, GA), and Cassia LLC (San Diego, CA). Fresh standards were made up to approximately 50 pmol/µL with ultrapure water with 0.1% piperidine. After analysis by the method of Swift Universal Glycan Acquisition (SUGA), as described^15^,, standard solutions were dried, and resuspended in ultrapure water with the dextran ladder.

### DDA Untargeted SUGA

Each sample was adjusted to 0.1% piperidine to enhance ionisation. The Thermo Tribrid Lumos was set to negative mode in DDA MS2 mode with a total cycle time of 6 seconds. Precursor spectra (540 – 1800 *m/z*) were collected in an Orbitrap at 120,000 resolution to hit an AGC target of 4e6. The maximum inject time was set to 600 ms. Fragment spectra were collected at 0.6 *m/z* resolution to hit an AGC target of 1e5 with a maximum inject time of 600 ms. The isolation width was set to 1.5 *m/z* and HCD normalised collision energy was set to 38. Only precursors charged between -1 and -6 were selected for MS2.

### Liquid Chromatography – Mass Spectrometry LC-MS setup for site A

Glycans were separated at site A with a ThermoFisher Scientific Vanquish Horizon HPLC and ionised into an Altis Plus triple quadrupole mass spectrometer. A single Hypercarb PGC column (100 mm length by 1 mm internal diameter, 3 micron pore size) was used for all separations. Both mobile phases contained 5 mM HFIP and 5 mM butylamine, with mobile phase A composed of water and mobile phase B composed of 40% water and 60% acetone. LC separation was performed at 155 µL/min.. For each injection, 10 μl was loaded and eluted using variable length gradients. The initial 40 min gradient was 0-37% mobile phase B over 30 min, held at 99% B for 5 min, then held at 0% B for 5 min. The final 80 min gradient used for glycans released from complex samples was 0-37% mobile phase B over 70 min, held at 99% B for 5 min, then held at 0% B for 5 min. Data were acquired using SRM.

### LC-MS Setup for Site B

At site B, glycans were separated with a Thermofisher Scientific Vanquish Horizon HPLC and ionised into an Altis triple quadrupole mass spectrometer. A self-packed^28^ PGC column with a Kasil frit (100 mm length by 100 µm internal diameter, packed with 3 micron pore size Hypercarb) was used for all separations. Both mobile phases contained 5 mM HFIP and 5 mM butylamine, with mobile phase A composed of water and mobile phase B composed of 20% water and 80% acetone. LC separation was performed at 40 µL/min. The LC gradient was 0-15% mobile phase B over 31 min, held at 99% B for 3.5 min, then held at 0% B for 5.5 min. Data were acquired using SRM.

### QQQ Method Development

A minimum of 13 of the most intense, monoisotopic product ions from the most intense MS2 spectrum acquired by SUGA were subjected to an SRM method for an automatically generated collision energy optimisation method in Skyline, at 9 CE steps from 10 to 90 V. In addition, the precursor *m/z*, and one fragment for each of the 11 dextran ladder standards were monitored. *N*-glycan mixtures, comprised of all available isomer standards, were created, acquired, and manually analysed to filter for quantitative (most intense) and qualitative (most discriminatory) product ions, resulting in a total of 3 product ions per structure.

### Applied Targeted Glycomics Method

429 unscheduled SRM transitions were selected, representing 183 mammalian *N*-glycan structure standards, with an additional 6 transitions, monitoring 6 dextran ladder subunits (3-8), to form the final mammalian *N*-glycan assay. For initial application of the method to bovine fetuin, human IgG, ovalbumin, and Xolair glycoprotein, a 40 min LC-MS method was used. For all other analyses, an 80 min LC-MS method was used.

## Data Analysis

*N*-glycan standard SUGA data was analysed by GlyCombo^29^ (v0.1) to identify glycan compositions, identify the most intense MS2 scans for each structure for annotation in GlycoWorkBench^30^, and construct the Skyline assay verifying compositional, and structural purities. Skyline^31,32^ (v24) was used to normalize RT, identify optimal CE for each product ion, create spectral libraries, quality filter identifications (>=0.9 idotp for SUGA, and >= 0.85 dotp for SRM analysis by spectral library), and quantify glycans. RT normalisation in the form of GU values were assigned to structures using the dextran ladder subunits 3-8 and a linear regression model. Spectral libraries were created from pure standard SRM analysis, and verified by applying the final SRM assay to isomeric mixtures of each standard. GlycoStore^17^ PGC RT values were retrieved on 2022/01/10 and are currently publicly available on GlyCosmos^33^ at https://glycosmos.org/glycostore/pgc. Comparisons between *N*-glycopedia were limited to fully defined structures shared between the two datasets. Glycan composition relationships were visualised with Compozitor^34^.

## Data Availability

The raw MS glycomics data generated in this study have been deposited in the GlycoPOST database^35^ under accession code https://glycopost.glycosmos.org/entry/GPST000578. All raw data, spectral libraries, and processed Skyline documents are available on Panorama^36^ (https://panoramaweb.org/Nglycopedia.url). Protocols for glycomics analysis are publicly available through the National Center for Functional Glycomics (https://research.bidmc.org/ncfg/protocols).

## Acknowledgements

The authors thank Jamie Heimburg-Molinaro for critical reading and manuscript editing, and Catherine A Hayes for providing access to GWS Parser which was used to rapidly retrieve GlyTouCan accessions from GlycoWorkBench. This research was facilitated by access to Sydney Mass Spectrometry, a core research facility at the University of Sydney.

The following reagents were produced under HHSN272201400008C and/or obtained through BEI Resources, NIAID, NIH: Spike Glycoprotein (Stabilized) from SARS-Related Coronavirus 2, Wuhan-Hu-1 with C-Terminal Histidine Tag, Recombinant from Baculovirus, NR-52308, Spike Glycoprotein (Stabilized) from SARS-Related Coronavirus 2, Wuhan-Hu-1 with C-Terminal Histidine and Twin-Strep^®^ Tags, Recombinant from HEK293 Cells, NR-53589, Spike Glycoprotein (Stabilized) from SARS-Related Coronavirus 2, Wuhan-Hu-1 with C-Terminal Histidine and Twin-Strep^®^ Tags, Recombinant from CHO Cells, NR-53937.

