## Supplementary Figures for "*N*-Glycopedia: Libraries for Native *N*-glycan Structural Analysis"

**Running Title:** A library of N-glycan standards enables targeted glycomics

**Table of Contents**

- **Figure S1** All unique *N*-glycan structure standards analysed to generate *N*-glycopedia and subsequently inform a targeted MS method
- **Figure S2** Compozitor output for the 91 glycan compositions assayed in *N*-glycopedia
- **Figure S3** Free *N*-glycan peak width is inversely correlated to PGC column temperature
- **Figure S4** Vendor cross-validation for six *N*-glycan structures confirms structural purity and identity
- **Figure S5** Unscheduled SRM enables quantitation of glycans not covered by *N*-glycopedia
- **Figure S6** MS2 matching scores across different mass spectrometers improve with recalibrated collision energy

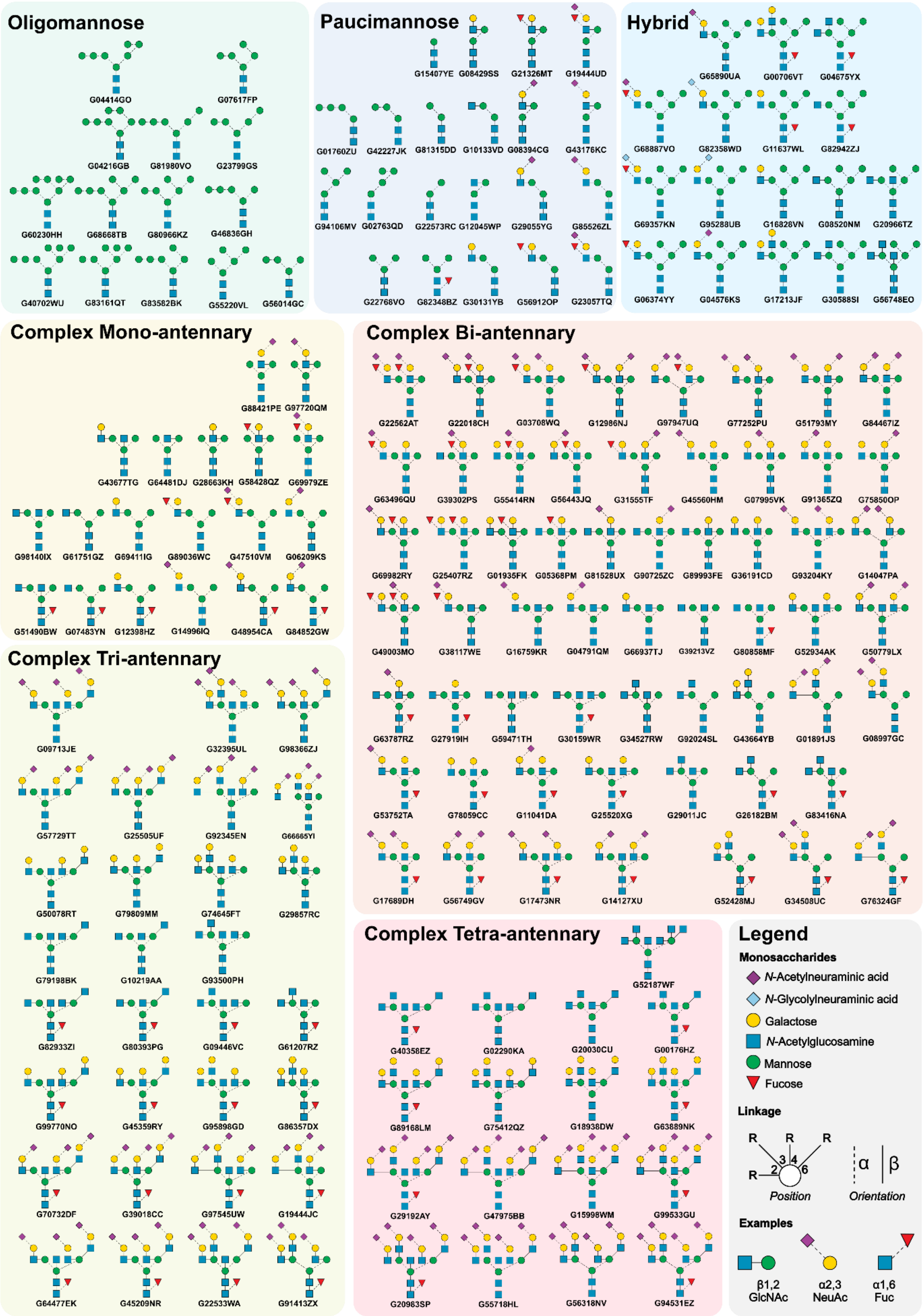


**Figure S1** All unique *N*-glycan structure standards analysed to generate *N*-glycopedia and subsequently inform a targeted MS method

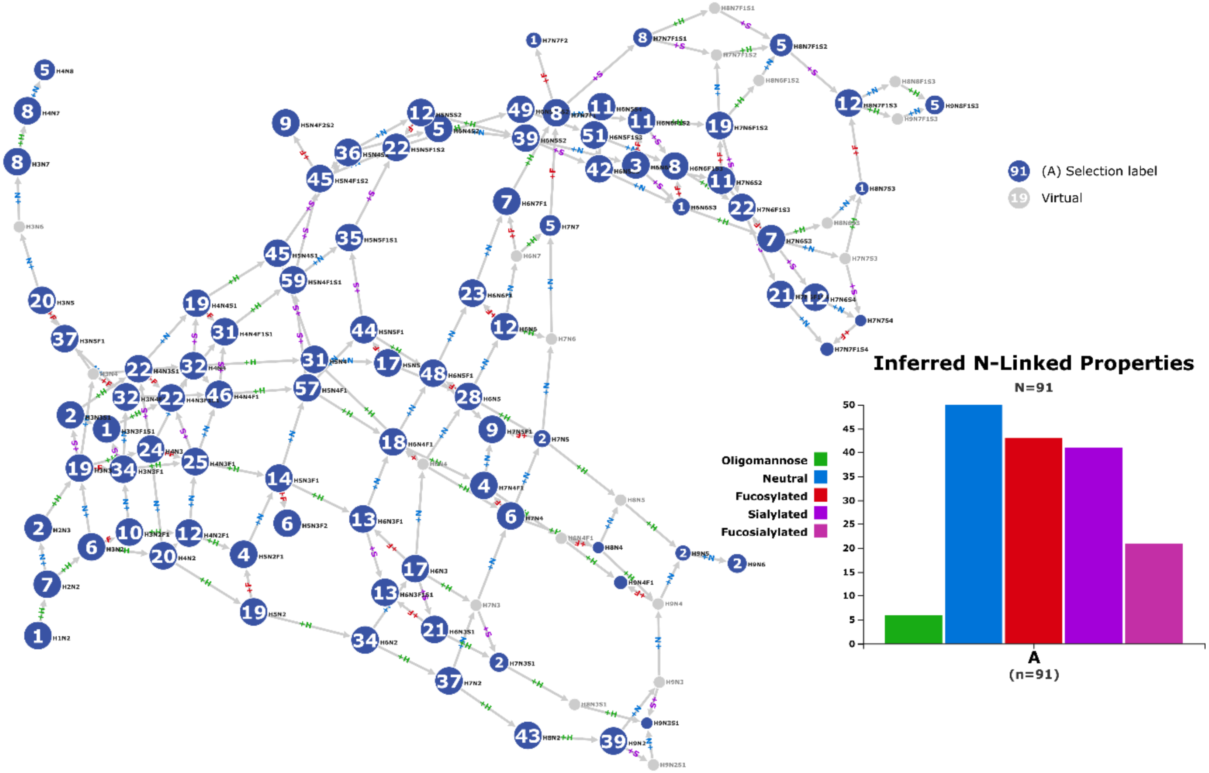


**Figure S2** Compozitor output for the 91 glycan compositions assayed in *N*-glycopedia

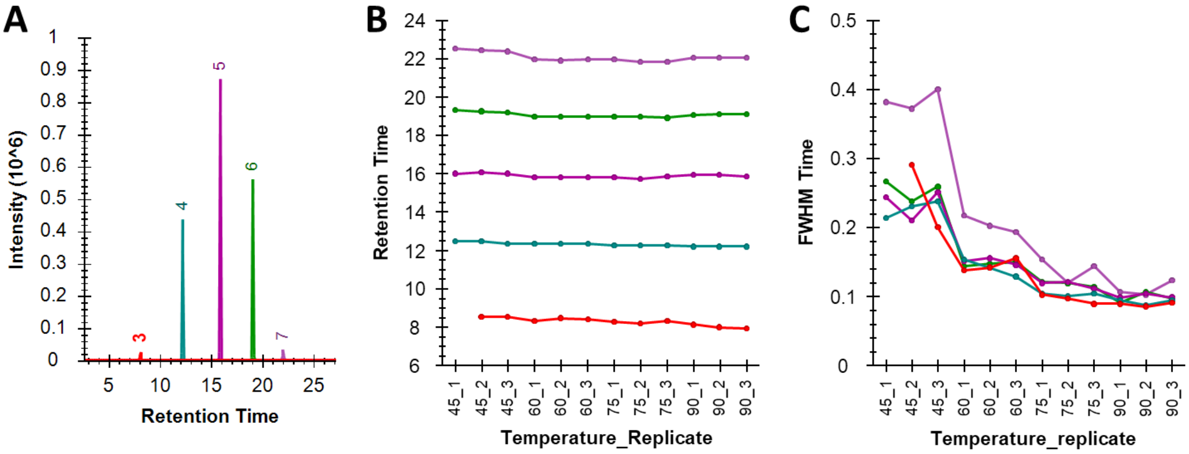


**Figure S3** Free *N*-glycan peak width is inversely correlated to PGC column temperature

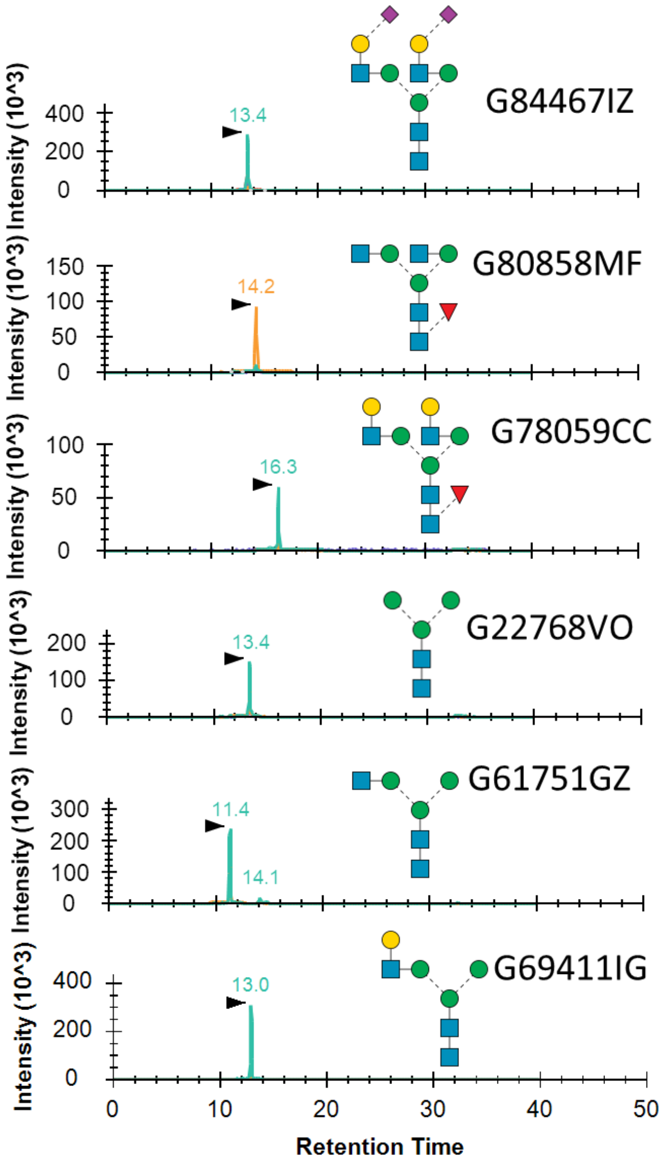


**Figure S4** Vendor cross-validation for six *N*-glycan structures confirms structural purity and identity

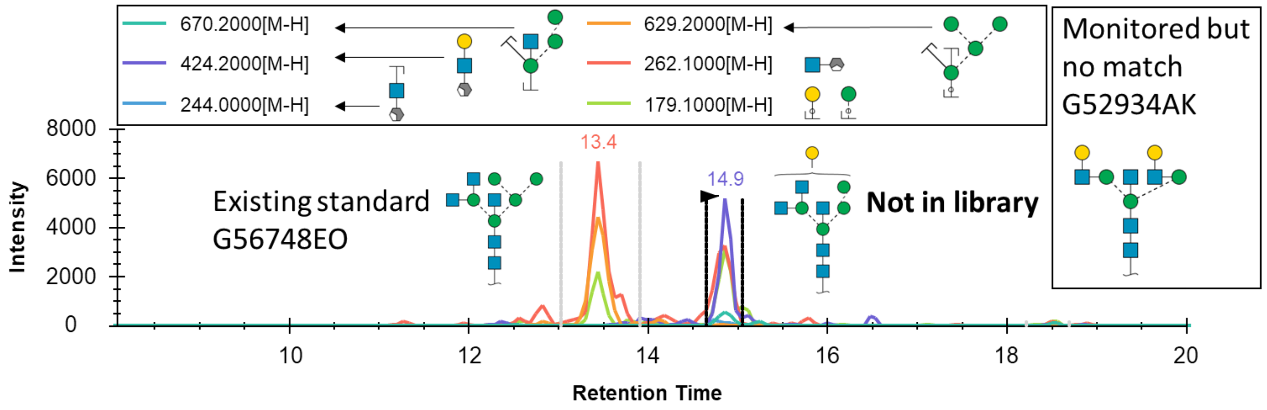


**Figure S5** Unscheduled SRM enables quantitation of glycans not covered by *N*-glycopedia

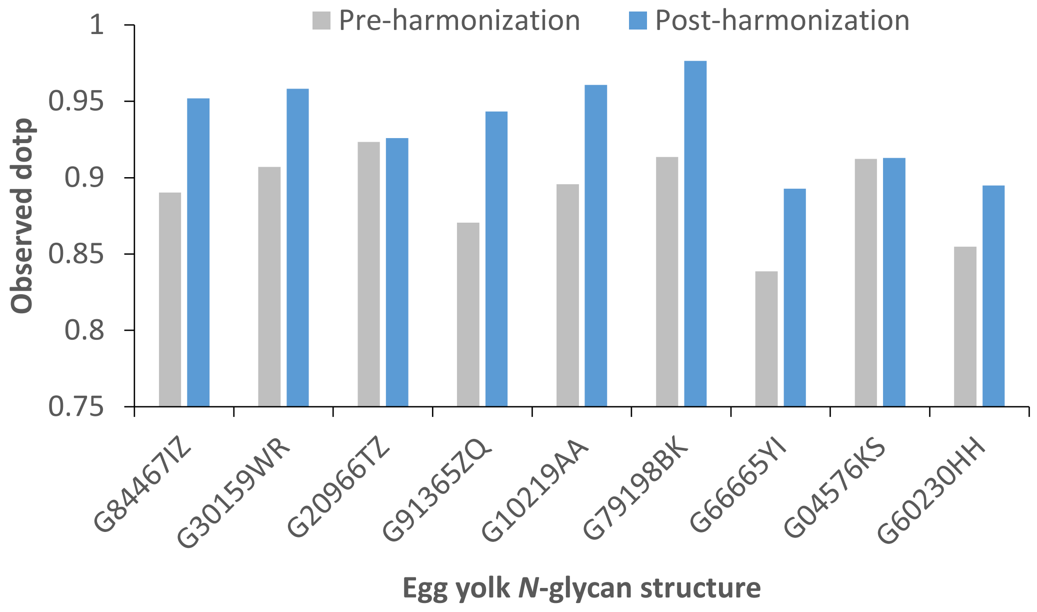


**Figure S6** MS2 matching scores across different mass spectrometers improve with recalibrated collision energy
